# Gray bats spatially segregate when navigating flight with conspecifics in complex environments

**DOI:** 10.64898/2025.12.26.696615

**Authors:** Eighdi Aung, Nicole Abaid

## Abstract

Bats are fast fliers with great maneuverability. Many species sense and communicate through echolocation, relying on acoustic signals in the environment. Bats also form large colonies, which necessitates that they fly in groups. When the group size increases or obstacles are in their flight paths, the acoustic space can become cluttered, posing a non-trivial challenge for navigation. How do they interact with their environment and conspecifics, and how do they balance social and environmental cues? Recent research has uncovered individual adaptations to calling and flight patterns when flying in the groups or avoiding obstacles. Studies also suggest coordination when foraging, as well as strategies used by the group to mitigate acoustic clutter. However, there has yet to be a study on how bats weigh different navigational tasks when flying in spatially complex environments. Here, we collected stereoscopic video data on a wild colony of gray bats, *Mytotis grisescens*, navigating their usual foraging flight paths in the presence of novel obstacles. We find that bats tend to stay close to a wall, but space out when flying in groups. We developed a data-informed agent-based model which revealed that their social repulsive behavior is strengthened when challenged to navigate novel obstacles.

## Introduction

Animals moving in the physical world must navigate complex environments, such as orienting themselves in response to chemical and flow gradients or avoiding collisions with obstacles. At the same time, social animals often perform coordinated motion, such as the striking patterns exhibited by starling flocks (1). The task of group navigation in the presence of environmental complexity requires an individual to actively attend to social cues while also dynamically responding to the heterogeneity in the space. This means that, for a group to navigate complex environments, its members must constantly strike a balance between social signals and environmental information. Failure to maintain this appropriate balance can result in inefficient or unfavorable outcomes for the individuals in the group. In the case of schooling fish, overreliance on social cues can reduce responsiveness to external stimuli (2), which in turn can increase energy expenditure and mortality. Conversely, overemphasis on environmental stimuli can cause fragmentation of the collective (3), which may increase predation risk in natural environments.

Balancing social and environmental cues may become more challenging when individuals are moving at high speeds and need to rely on noise-prone sensory modalities such as echolocation. Bats (order *Chiroptera*) navigate these challenges well, being the only mammals that are capable of flapping flight with speed and agility. For example, Brazilian free-tailed bats (*Tadarida brasiliensis*) can reach speeds up to 30 m/s (4), and lesser horseshoe bat (*Rhinolophus hipposideros*) are known to be able to make 89.5° turns with lateral accelerations of 17.5 m/s^2^ (5). Many species of microchiropteran bats are known to primarily rely on echolocation to sense their surroundings when navigating (6). Unlike vision, echolocation is an active form of sensing where a bat sends out acoustic signals and listens to the reflected echoes to sense their surroundings. As social animals, they often fly in groups, which can complicate their navigation, particularly because of the potential overlap of echoes in the environment as the density of the swarm increases (7–10). This challenge in group navigation can become more complex when flying in cluttered environments where the echoes can reflect off many objects in the space (11). In obstacle-ridden environments, bats need to increase the number of echolocation calls they make to sense obstacles in their flight path while moving at fast speeds (12), but at the same time overcome the cocktailparty problem of acoustic signals. This poses a non-trivial decision making problem that begs the question: how do bats balance these competing cues?

How bats can navigate, respond to obstacles, and adapt socially has been well studied from a behavioral perspective. Bats are known to use spatial memory for foraging and navigation, relying on both externally observed landmarks (13, 14) and internally stored representations of the space (15–17) to fly through familiar terrain. Even when their learned environment changes, bats are still able to adapt their individual behaviors to avoid obstacles. For example, experiments with big brown bats (*Eptesicus fuscus*) revealed that, when vertical gaps between obstacles narrow, they fly slower, higher, and emit grouped bursts of echolocation pulses with shorter intervals to navigate the gaps (12). Greater horseshoe bats (*Rhinolophus ferrumequinum nippon*) are also known to gradually reduce their meandering, narrow their pulse-direction shifts, and emit fewer but more precisely aimed calls as they learn the layout of the new environment (18). When species of bats that heavily rely on echolocation for navigation fly together, they may call louder and longer (19) or shift their calling frequencies (8) to overcome signal jamming. Additionally, there are also studies on pairwise social interaction in bats, suggesting rear-to-front information transfer (20, 21) or the presence of delayed leader-follower heading matching (22, 23). A more recent study also suggests that they may spread out spatially to possibly overcome acoustic clutter (24).

While recent studies have revealed bats’ biological capabilities and behavioral adaptations to environmental and social cues, it remains an open question how a bat weighs these different cues. Even more so, only a limited body of work has attempted to model the mechanism of the response to the environment and to conspecifics. Hallam et al. (25) modeled the column-like patterns of Brazilian free-tailed bat (*Tadarida brasiliensis*) swarms using flocking, repulsion, and predation avoidance mechanisms. Giuggioli et al. (26) modeled the chasing flight patterns of foraging Daubenton’s bats (*Myotis daubentonii*) in the form of delayed flocking. Lin et al. (27) modeled bats’ goal-directed and avoidance behaviors from modeling the response of reflected echoes from the environment. More recently, Vanderelst (28) modeled the collective movement of bat swarms following acoustic gradients for obstacle avoidance. From those limited models, only Lin’s and Vanderelst’s account for responses to both environment and social forces. However, because Lin collapsed the two responses into a single avoidance force, the balance of cues cannot be determined. Likewise, Vanderelst did not differentiate acoustic responses between the environment and conspecific, thereby also limiting the ability to evaluate the relative weighting of these cues.

In this paper, we seek to address two research questions: how do bats interact with their environment and with their conspecifics, and how do they balance social interactions against interactions with the environment? We answer these questions through an experimental study on a wild gray bat (*Myotis grisescens*) colony with introduced environmental obstacles. We statistically compare the behaviors of bats with and without obstacles as assessed through video data. Informed by our statistical analysis, we introduce an agent-based model grounded in simple interaction rules, and validate our model with experimental data to discover how bats weigh navigational cues from social forces and the geometry of their environment.

## Data collection and analysis

### A. Experimental data

The data were collected on a bridge over Beaver Creek in Bristol, Virginia, USA in 2022, from a maternal colony of approximately 10,000 gray bats, *Myotis grisescens*. During the summer season, the bats exit a culvert on the creek nightly to hunt for insects, starting at approximately ten minutes after sunset. They fly upstream towards the bridge, where the recording equipment is located. We used 4 GoPro cameras (Hero 3+, Black Edition) modified with infrared (IR) sensitive lenses (IR-Pro IRP202 Hybrid lenses), which allow the cameras to be sensitive to infrared light. The cameras recording at 60 frames per second, imaging a volume of approximately 14 m cross stream, 6 m upstream and 3 m deep. Since the bats fly out at dusk, we used five infrared lights (Tendelux, CCTV Lighting, Mid-Long Range IR Illuminator) to illuminate the space. Data were collected for two nights on the 23rd and 24th September of 2022 for approximately 45 minutes each night, starting around 20 minutes before sunset. On the first night, the bats were allowed to fly in their natural environment. On the second night, two obstacle arrays made out of foam pool noodles, strung across the creek on nylon ropes, were used to introduce novel obstacles to the bats’ flight paths. Figure 1 shows the data collection site and the experimental setup.

**Fig. 1.**
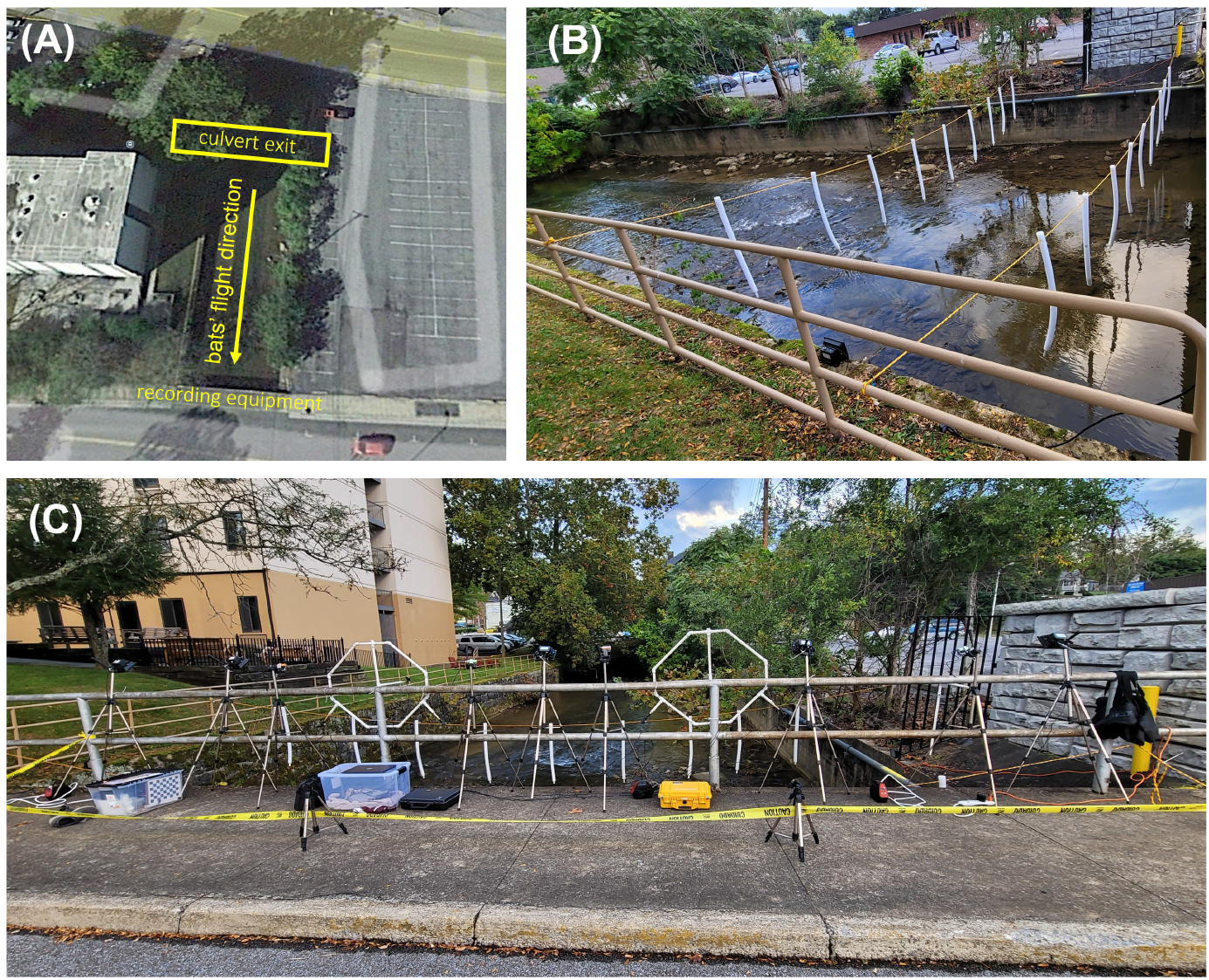
Experimental setup: (A) Google map view of experimental site at 36°35’52.6”N 82°10’57.0”W. Bats fly out of the culvert exit at sunset and upstream towards the bridge where the recording equipment is set up. (B) The obstacle arrays comprising 18 pool noodles strung in staggered positions using 2 nylon ropes. (C) Recording equipment set up on the bridge, consisting of GoPro cameras, ultrasonic microphones, and IR lights.

The GoPro cameras had lens distortion at the edges, which was corrected by using a custom Matlab script. From the corrected video frames, we calibrated the cameras for stereoscopic tracking. We first estimated the intrinsic camera parameters using MATLAB’s built-in CameraCalibrator function (29), and the extrinsic parameters using EasyWand5 software (30). To compute the intrinsic camera parameters, specifically the focal length and principal points of the of each of the four cameras, we used a checkerboard with 25mm checks. Each camera was calibrated by recording a minute-and-half video where the checkerboard is moved in the view of the camera at different angles. We then calibrated for extrinsic camera parameters, specifically the three angles of 3D rotation and three parameters of 3D translation, which required a procedure of moving a cross-shaped wooden wand in the view of the cameras. We followed the detailed procedure provided by the EasyWand5 software (30) for these calibrations.

Using the coefficients obtained from the EasyWand5 software, we imported the coefficients into the DLTdv8 software (31) to perform 3D tracking of the flying bats. For both nights of the experiment, we tracked three ten-second clips spaced 20 seconds apart. The 3D points obtained from DLTdv8 have an arbitrary reference frame that can change across different calibration datasets, therefore the tracked points between two nights are not in the same reference frame. To allow for direct comparison and analysis between the two experiments conducted at different nights, we used the points on a physical axis marker to rotate the 3D points to co-local reference frame, and used a common feature in all experimental trials to translate to the same world coordinates. After obtaining 3D points in identical reference frames, we smoothed the bat trajectories using a moving average over window of 5 video frames (0.083 s). Additionally, we translated and reflected the 3D points such that the bats’ flight direction upstream is along the positive x axis and the cross stream direction is along the positive y axis. Figure 2 shows the data processing pipeline.

**Fig. 2.**
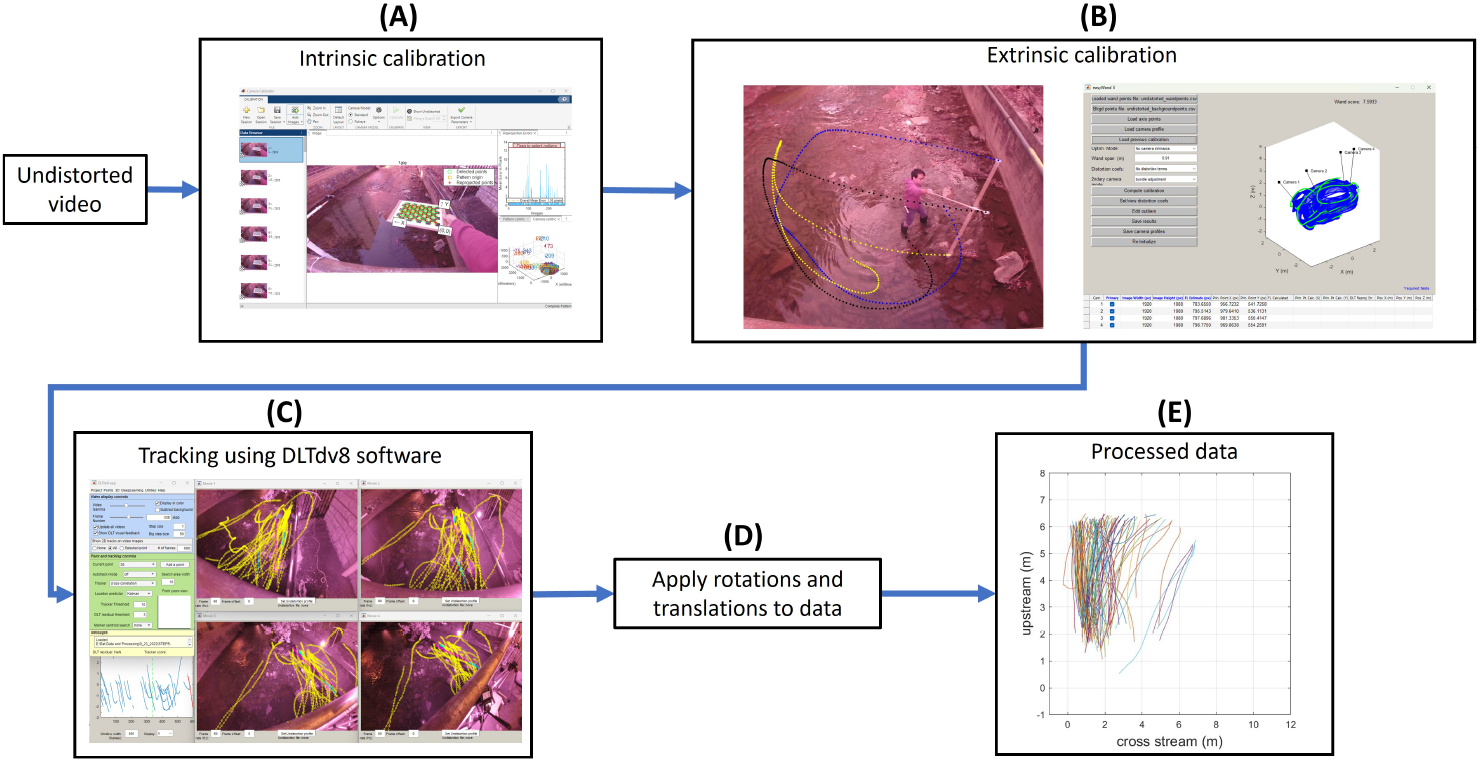
Data processing pipeline: (A) intrinsic calibration using MATLAB’s built-in CameraCalibrator function (29) to obtain camera focal length and principal points; (B) extrinsic calibration using EasyWand5 software (30) to obtain rotation and translation matrices that define the camera’s position and orientation in the world coordinates; (C) stereoscopic 3D reconstruction of 2D camera data using the DLTdv8 software (31); (D) 3D data rotated and translated to align and match the coordinate systems of with and without obstacles; (E) processed data used for analysis and modeling.

A single bat takes approximately one second to fly through the imaged volume. Over the span of the two nights, we tracked 37 single bats flying in the absence of obstacles, and 41 single bats flying with obstacles, whose trajectories are used for identifying individual dynamics. From three 30-second clips for each night, when bats are flying in groups, we tracked 121 bats for the night without obstacles, and 100 bats for the night with obstacles. To estimate the positions for the obstacles in the space, we tracked the top and bottom points of the obstacles. Using the known diameter of the pool noodles, we estimated the obstacles as cylinders. To orient the axis in each view and to resolve the coordinates between experiments, we also tracked the tips of a 3D axis marker object made of PVC pipes, and a boulder in the middle of the creek for a reference axis and a world point, respectively.

### B. Preliminary behavioral analysis

The overall dynamical behavior of bats can be summarized by the distribution of their time-averaged velocities. Figure 3 shows violin plots of average velocity of bats in the upstream direction, *V*_*x*_, the cross stream direction, *V*_*y*_, and the direction of gravity, *V*_*z*_. Due to the non-normality and heteroscedasticity of the distributions, we select the Mann-Whitney U-test to statistically compare datasets. In the interest of modeling bats’ group motion, we test for significance of the alternative hypothesis that the bats’ velocities (when in groups) are different between the two experimental conditions defined by the presence and absence of obstacles. We find significant differences in the velocities in the upstream direction (*W* = 8008, *p* < 0.001) and the cross stream direction (*W* = 2981, *p* < 0.001), but no significant difference in the direction of gravity (*W* = 4534, *p* = 0.46). This results suggests the bats’ responses to the experimental obstacles are dominantly planar. Informed by this result, we chose to reduce our study of the experimental data and the resulting model to two dimensions, focusing only on the behavior of bats in the upstream and cross stream directions.

**Fig. 3.**
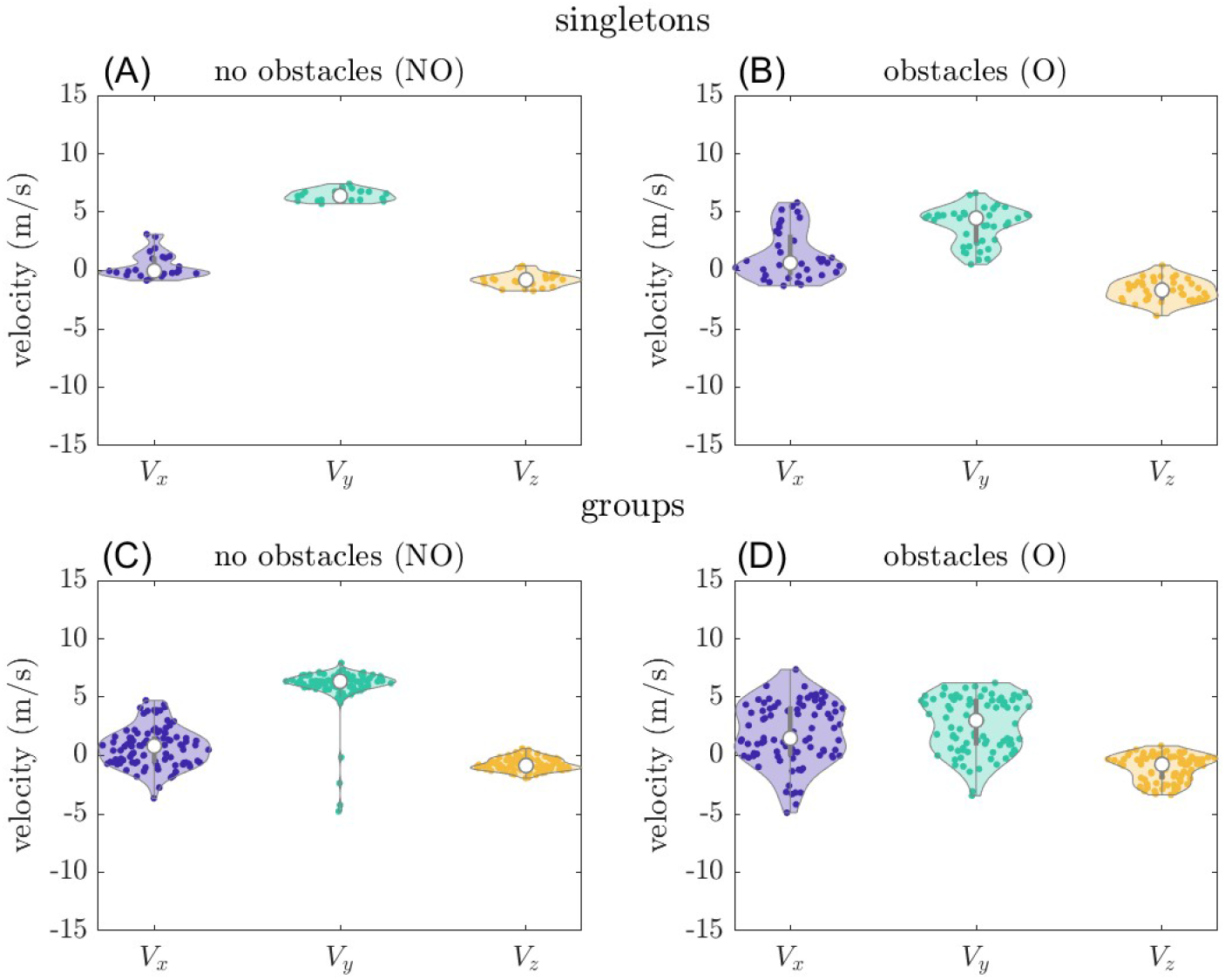
Bat flight velocities: Violin plots showing the distribution of average upstream velocity, *V*_*x*_, cross stream velocity, *V*_*y*_, and velocity in the direction of gravity, *V*_*z*_ for (A) singleton bats without obstacles, (B) singleton bats with obstacles, (C) groups without obstacles, and (D) groups with obstacles. The violin plots are generated using an open-source MATLAB software by Bechtold (32).

Furthermore, we examine the effect of sociality on the flying behavior of bats. With the acquired dataset on singleton bats, we test the hypothesis that the bats’ velocities are different when flying in groups. For the case of no obstacles, we do not find significant differences in the velocities both upstream (*W* = 982, *p* = 0.34) or cross stream (*W* = 1213, *p* = 0.57). For the case with obstacles, we find significant differences in the velocities for upstream (*W* = 1278, *p* = 0.03), but not for cross stream (*W* = 2001, *p* = 0.11). These results suggest that flying in groups significantly effects the bats’ upstream velocities when obstacles are present. Qualitatively, Figure 3 also shows that, when obstacles are present and bats are flying in groups, there is a larger variance in the velocity of bats in the 2D plane of interest.

Figure 4 shows the spatial trajectories of bats flying with and without obstacles from a top-down view of creek. The multicolored lines mark the positions of bats and the red circles show the location of obstacles. Considering the distributions of the bats’ positions upstream and cross stream over the tracked 30 seconds, we found that the bats mostly fly straight and prefer the left side of the creek when there are no obstacles, see Figure 4 (A) and (C). The zero cross stream axis runs along a wall parallel to the creek. The behavior of bats choosing to fly around a meter distance from this wall may indicate a thigmotaxis or wall-hugging behavior, a behavior where organisms tend to stay close to linear structures as a guide for navigation. Additionally, comparing Figure 4 (A) and (C), we can qualitatively see that bats fly further across the stream when flying in groups. We test the hypothesis that bats’ time-averaged positions across the stream, in the absence of obstacles, differ between solitary and group flight. When flying alone, bats fly at an average position of 1.2 m, and 1.8 m when flying in groups. We find significant differences in the mean and variance in cross stream positions between singleton and group flight when applying the Mann-Whitney U-test (*W* = 1721, *p* < 0.001) and Levene’s test (*F* = 5.27, *p* < 0.05). We also see similar behavior in singleton bats when obstacles are present, Figure 4 (B). A subset of bats exhibit a strikingly different flight path, turning away from the wall and flying perpendicular to the stream, Figure 4 (B) and (D). The difference in the spatial distribution of bats between the experimental conditions qualitatively supports the statistical significance that we find in the bats’ velocities, suggesting that bats are dynamically responding to the obstacles by turning across the stream.

**Fig. 4.**
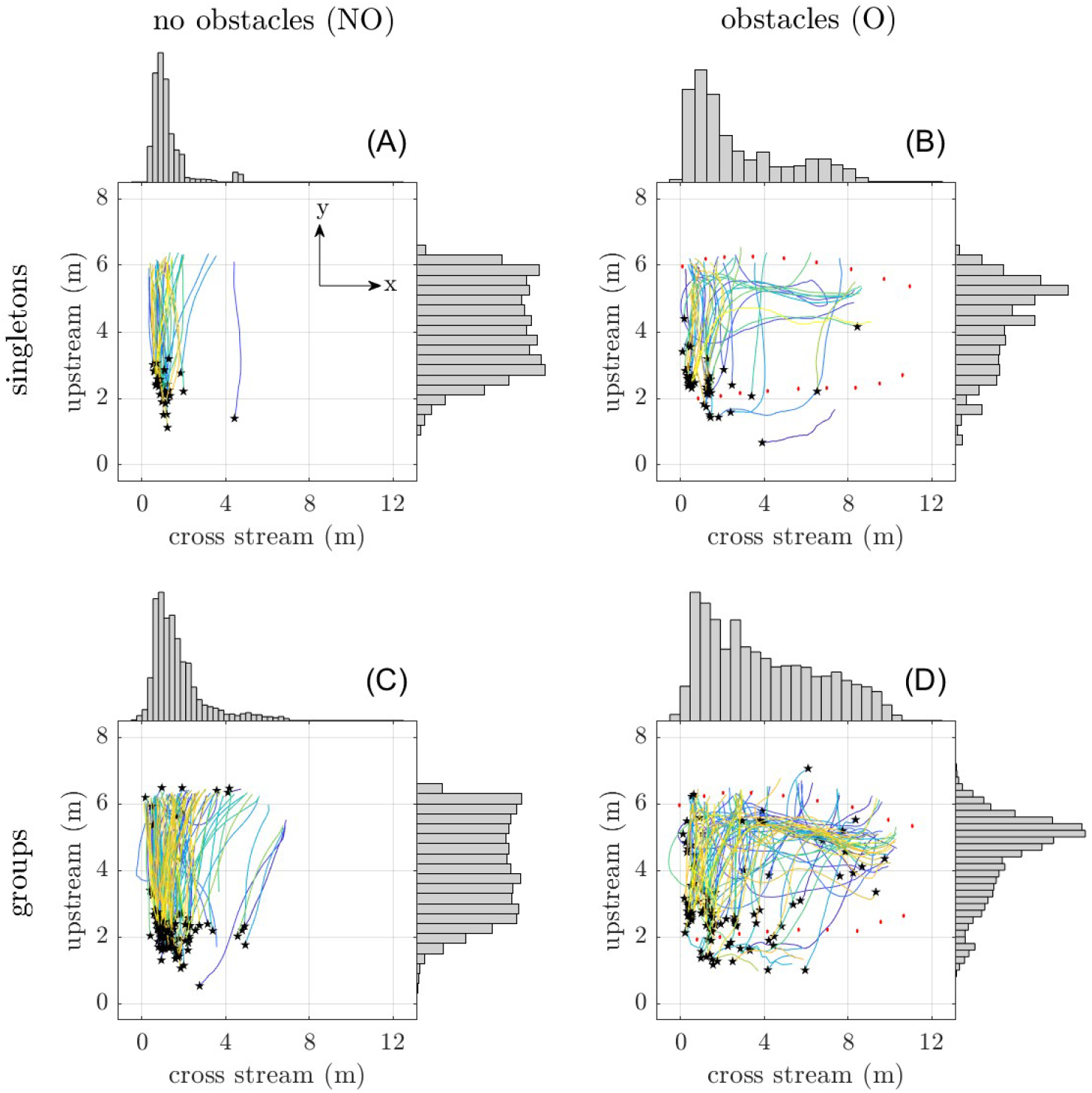
2D trajectories of all tracked bats: (A) singleton bats without obstacles, (B) singleton bats with obstacles, (C) groups without obstacles, and (D) groups with obstacles. Each of the colored lines shows the trajectory of a unique bat, where the black star marks the start of the trajectory. The histograms on the top of each subplot correspond to the distribution of all the cross stream trajectory points, and histograms on the right of each subplot correspond to the distribution of all the upstream trajectory points. Red circles in the right figures show the locations of the experimental obstacles’ centers.

## Modeling and simulation

We model a group of *N* agents, bats, navigating a 2D channel spanned by two vertical walls, simulating the two experimental conditions with and without obstacles. In the simulations, the x-axis and y-axis correspond to the cross stream and upstream directions, respectively.

### C. Social interaction rules

We can infer the prevalence of social interaction rules that the bats exhibit from the data by applying fundamental principles of collective behaviors – namely, attraction, repulsion and flocking (33). Attractive or repulsive interactions can be modeled as an adjustment in the relative distance between focal and the nearest neighbor bat. Flocking can be modeled as an adjustment in the heading of a focal bat relative to the heading of its’ neighbor. We applied linear mixed effect models detailed in Supplementary Section 1, and Figure 5 shows the fits obtained from the models. The slopes obtained from the models inform the strength of attraction/repulsion and flocking. We obtain the slope for attraction and repulsion to be *β*_1_ = 0.15, *p* < 0.01, and *β*_1_ = 0.24, *p* < 0.01 in the absence and presence of obstacles, respectively. For the flocking interaction, we obtain slopes *β*_1_ = 0.070, *p* < 0.01, and *β*_1_ = 0.0087, *p* = 0.098 in the absence and presence of obstacles, respectively. This result shows that attraction or repulsive interactions dominate collective motion in both experimental conditions as quantified by the magnitude of the fit slopes, and that flocking motion is not significant when obstacles are present.

**Fig. 5.**
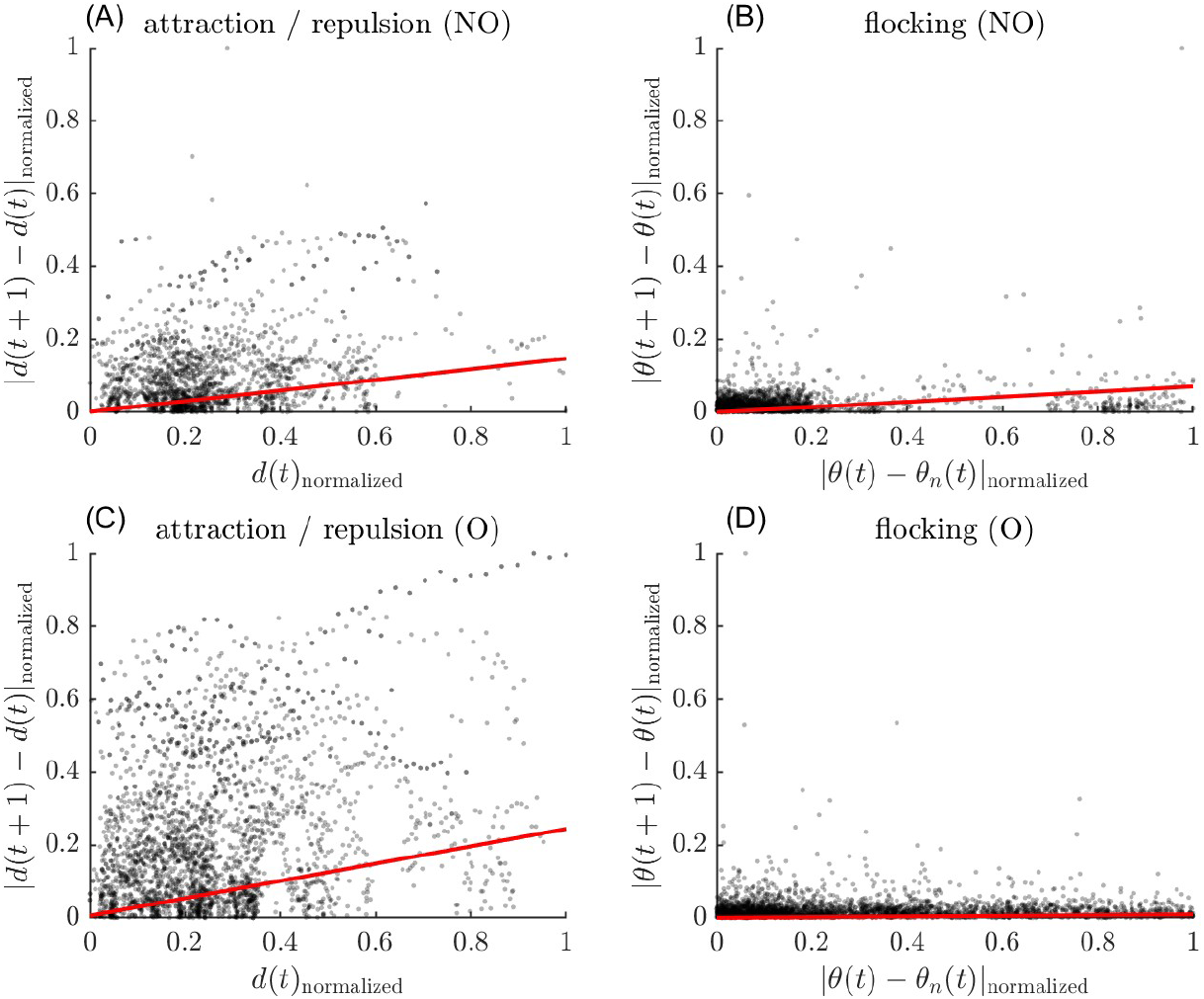
Analysis of interaction rules using linear mixed models: (A) attraction/repulsion rule without obstacles, (B) flocking rule without obstacles, (C) attraction/repulsion rule with obstacles, and (D) flocking rule with obstacles. Here, *d*(*t*) represents the distance to the nearest neighbor, *θ*(*t*) represents the heading of the focal bat, and *θ*_*n*_(*t*) represents the heading of the nearest neighbor bat at time *t*. A forward-facing model is also imposed, i.e., a bat is a nearest neighbor to a focal bat if and only if it is also in front of the focal bat with respect to its velocity vector. The linear fits are shown in red, and the slopes, *β*_1_, and p-values are: (A) *β*_1_ = 0.15, *p* < 0.01 (B) *β*_1_ = 0.070, *p* < 0.01 (C) *β*_1_ = 0.24, *p* < 0.01, (D) *β*_1_ = 0.0087, *p* = 0.098.

### D. Agent-based modeling

Informed by the analysis of bats’ velocities and spatial distributions, we propose a simple agent-based model that focuses on three behavior rules: (1) channel-following behavior; (2) social behavior in the form of repulsive interactions with neighbors; and (3) avoidance of obstacles. In this model, we assume that the agents only respond to objects or other agents in front of them, which is a behavior of bats when first exiting a hibernaculum (23, 34). Let **x**(*t*) ∈ ℝ^2^ be the position of each agent at time *t*. Agents move at constant speed, *s*, with heading, *θ*(*t*) ∈ [− *π, π*], and velocity, **v**(*t*) ∈ ℝ^2^. At each time step, we update the velocity of an agent via a weighted sum of the inertia and the three behavior rules:

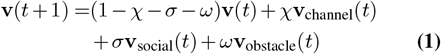

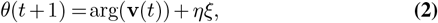

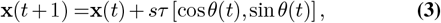

where *χ* is the weight for channel-following behavior, *σ* is the weight for social repulsion to nearest neighbors and, *ω* is the weight for obstacle avoidance, with *χ, σ, ω* ∈ [0, 1]. For modeling the condition where obstacles are absent, *ω* = 0. *τ* is the incremental time *s* is the speed, and *ξ* ∈ 𝒰 [0, *π*/2] is the angular stochastic noise with uniform distribution. After the positional update from behavior rules, we impose area exclusion by checking if agents are overlapping with any other objects in the space. If an overlap were to occur, i.e., the distance between the object and the agent is less than half a wingspan, the prior positional update is ignored, and instead the agent is turned away from the object:

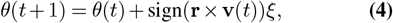

where **r** is the vector from the agent that is orthogonal to the object it would overlap with, and *ξ* ∈ 𝒰 [0, *π/*2]. If the agent were to violate two or more objects at once, the agent takes the mean heading obtained from each of the maneuvers. We also want to note that the violation events occur rarely in the simulations due to our parameter choices, i.e., < 5% of the time.

Below we detail how each of the behavior rules are implemented to compute the time evolution of bat velocities.

#### D.1. Channel-following behavior

Let *w* ∈ {left, right} represent the left and the right walls of the channel, and *x*_*w*_ represent the location of the walls in the cross-stream direction, i.e., *x*_left_ = 0 and *x*_right_ = 14. Let *R*_*w*_ be the distance at which bats prefer to stay from either wall. Then, the attractive forcing towards this distance, *R*_*w*_, from the walls can be modeled as *d*_left_(*t*) = |(*x*_left_ + *R*_*w*_) − *x*(*t*)| and *d*_right_(*t*) = |(*x*_right_ *R*_*w*_) − *x*(*t*)|. The velocity in response to the channel following behavior is computed by:

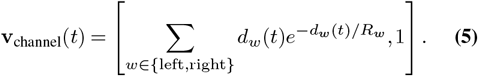

By construction of this velocity field, we allow the attractive force towards *R*_*w*_ to exponentially decay over distance. The resultant behavior replicates a goal-like forcing where agents align with upstream direction to follow the channel.

#### D.2. Social repulsion and obstacle avoidance behaviors

The social repulsion and obstacle avoidance are modeled in the same way as a repulsive interaction from an object of interest. To allow for a simple parameterization of this behavior, we define these repulsive interactions only to the closest object, i.e., the nearest neighbor or obstacle. The velocities **v**_social_ and **v**_obstacle_ are modeled by imposing an agent-based rule on the heading of the agents, *θ*_repel_. The heading in response to a repulsive behavior is computed by:

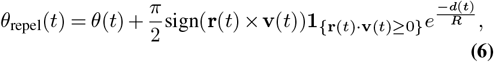

where **r**(*t*) is the vector from the agent that is orthogonal to the object the agent is avoiding which has length *d*(*t*), and *R* characterizes the length where the turning behavior lessens over distance. We will define *R* = *R*_*s*_ to characterize the social interaction, and *R* = *R*_*o*_ to characterize the interaction with obstacles. **1**_{}_ is an indicator function that returns a 1 if **r** · **v**(*t*) ≥ 0, which occurs only if the object of interest is in front of the agent and the turning is activated.

### E. Simulation conditions

The geometry of the environment is fixed for all simulations and is scaled to match the imaged area of the data. We define the walls to be parallel to the y-axis and to be positioned at *x* = 0 and *x* = 14, creating a channel of width of 14 m. In the environment with obstacles, we simulated two rows of nine obstacles each. The obstacles have radius of 0.0615 m, and are located where they appeared in the experimental data. Bats are simplified as circular agents with diameter equal to the average wingspan, *w* = 0.28 m. We impose an exclusion of area for each bat; therefore, each agent is prohibited from overlapping with walls, obstacles, or other agents in the space.

Agents are spawned in the upstream positions sampled from a uniform distribution, i.e., *y* ∈ 𝒰 [− 6, −2]. The spawning location for cross-stream positions are sampled from an empirical distribution estimated from experimental data of bats flying in groups without obstacles. Specifically, we extracted cross stream position data from four narrow horizontal bands centered at *y* = {2.0, 2.5, 3.0, 3.5}, each spanning ±0.1 m in the vertical direction. These data are used to construct a distribution from which the cross stream positions of simulated agents are drawn.

For fair comparison to the experimental data, the simulated trajectories are truncated to only consider the imaged area we record in the field experiments. This control area is [2, 6] m in the upstream coordinate and [0, 10] m in the cross stream coordinate. Additionally, we could not image the area before the bats first encounter the first row of obstacles, where they may be turning and entering the image area further down the cross stream direction. To account for this, we also only consider trajectories that enter the control area from *x* < 6. Any trajectory outside the control area or which does not satisfy the entrance criterion is not compared against the experimental data. See example simulated trajectories in Figure 8, where trajectories that satisfy the criteria are represented with darker lines and those that do not satisfy are translucent.

The number of the agents in the area must be allowed to varying with time throughout the simulation since the experimental data show that bats continuously flow in and out of the imaged area. To simulate and account for the non-constant number of agents in the system at any time, we control the flow rate of agents into the simulated area. The distribution of time lag between bats entering the control volume is expected and shown to follow an exponential distribution experimentally. We empirically estimate this distribution which is parameterized by a single parameter, *μ*, that corresponds to the mean number of frames of between the entrance of any two consecutive bats into the image area and use this distribution to generate initialize simulated trajectories.

**Table 1.** Summary of model parameters for conditions without obstacles (NO) and with obstacles (O), where *χ, σ*, and *ω* weight the channel-following, social repulsion, and obstacle avoidance behaviors, respectively.

| | $\chi$ | $\sigma$ | $\omega$ |
| --- | --- | --- | --- |
| NO | 0.035 | 0.004 | - |
| O | 0.009 | 0.015 | 0.2 |

### F. Model parameters

The model parameters are obtained from the experimental data. The speed of bats, *s*, is 7 m/s without obstacles and 5 m/s with obstacles. The time step for dynamic update, *τ*, is 1/60 s. The rate at which bats come into the simulation domain is every *μ* = 0.24 s. The distance, *R*_*w*_, at which the bats like to stay close to a wall is 0.8 m. The minimum distance at which bats fly with their nearest neighbors, *R*_*s*_, is 1.1 m. The minimum distance that bats fly near an obstacle, *R*_*o*_, is 0.5 m. The stochastic noise, *η*, is 0.05. Please see Supplementary Section 2 for details on parametrization.

## Model fitting for behavioral weights

The parameters that weight behaviors in the velocity update, *χ, σ*, and *ω*, are found via parameter sweep for the optimal values. For each unique set of parameters, we compute a metric from the simulated trajectories and compare its distribution to that of the same metric computed from the experimental data. Prior statistical analysis, using the Levene’s test, suggested that bats spread out cross stream due to flying in groups. Since the spread is most prominent in the cross stream direction, we define one of the comparison metrics to be the distribution of the mean cross stream positions of the trajectories, defined as 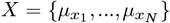, where 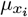 is time-averaged cross-stream position of bat *i*. Another notable quality we see in the behavior of bats in the presence of obstacles is turning in the cross stream direction. This can be represented by the distributions of velocity in the upstream direction since we also found statistical significance for differences in the upstream velocity when obstacles are present. We define this distribution as 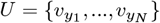, where 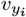 is the time-average cross-stream velocity of bat *i*. To quantify the sameness of distributions, we compute the energy distance defined by:

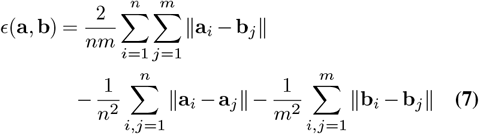

where **a** is the vector of *n* samples of the computed metric obtained from the simulations, **b** is the vector of *m* samples of the computed metric obtained from the experimental data, **a**_*i*_ and **b**_*i*_ are their elements. For the case with no obstacles, we minimize the energy distance using the metrics *X* and *U*. The heatmaps of the resulting energy distances are shown in Figure 6, obtained from a parameter sweep for *χ, σ* ∈ [0, 0.2] with increments of 0.001. Additionally, we also ran simulations with sparser increments of 0.01 to obtain highest and lowest possible energy distance for each metric in the full parameter space, *χ, σ* ∈ [0, 1], which we use to normalize the heatmaps. For each metric, locating the energy distance of *ϵ* = 0, in the normalized parameter space should yield a parameter set that best fits the data. Ideally, the optimal parameter value for one metric should also be optimal for the other. However, this requirement is overly conservative. To allow for a more realistic comparison, we introduce a tolerance, *δ* > 0, for which a set of parameter values can still be considered optimal across all metrics. We define this tolerance as the smallest value at which the first intersection of optimal regions across all metrics will be possible in parameter space. This translates to looking for regions in the heatmaps where the regions with energy distance less than *δ* for each metric will intersect in the parameter space.

**Fig. 6.**
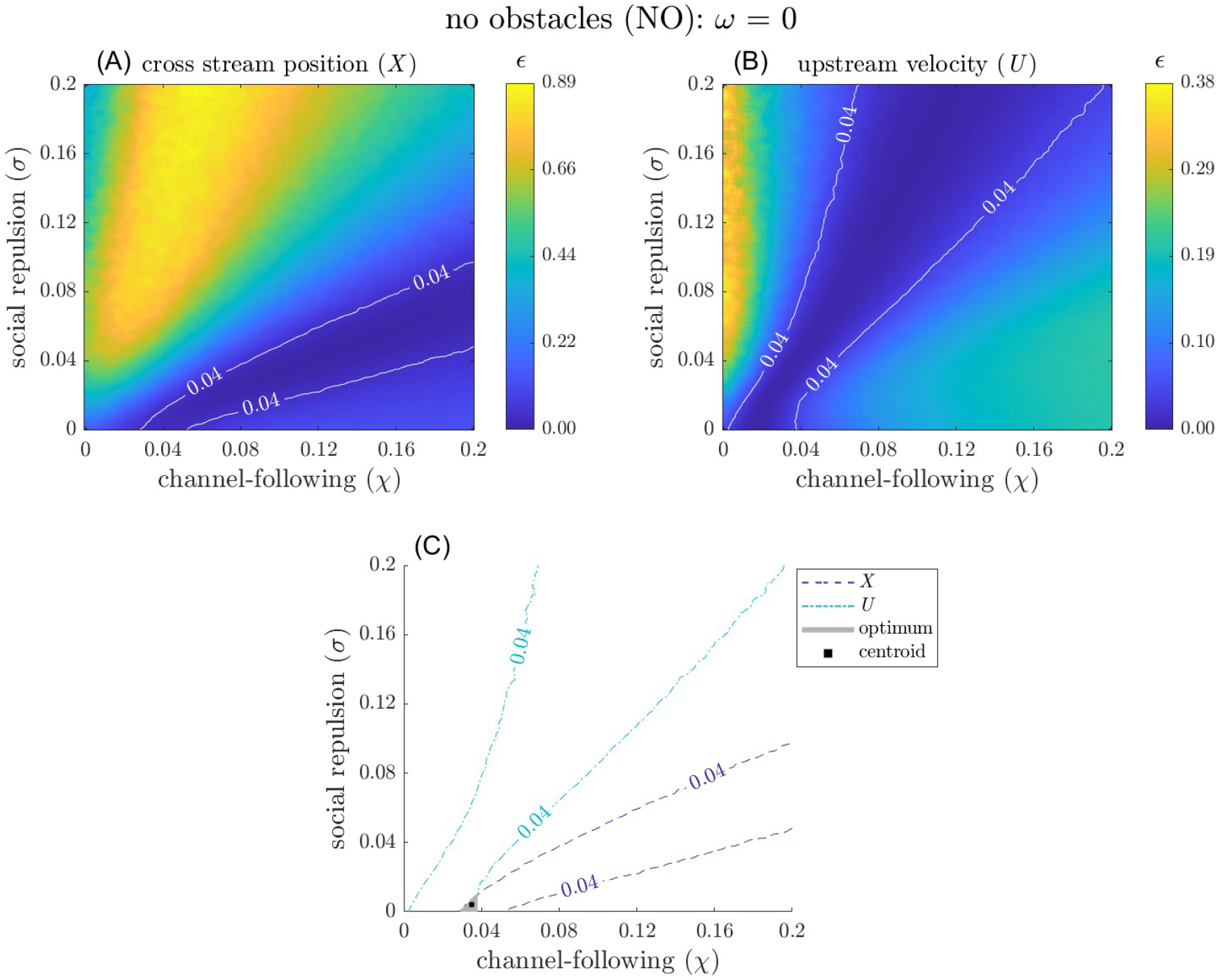
Heatmaps display values of energy distance, *ϵ*, computed using distributions of (A) cross stream positions and (B) upstream velocity. The color bars for each of the heatmaps represent the energy distances that are min-max normalized over the entire range of possible parameter values, i.e., *σ, χ* ∈ [0, 1]. The heatmaps shown only consider the extent at which the optimal value is found, i.e., *ω* = 0 for obstacle avoidance, *χ* ∈ [0, 0.2] for channel-following, and *σ* ∈ [0, 0.2] for social repulsion. White lines overlaid top of on each heatmap represent the error threshold, *λ* = 0.04. (C) Energy distance contours with intersection in the parameter space depicted by the shaded gray region. The centroid of the optimal parameter set is shown by a black square at *χ* = 0.035 and *σ* = 0.004.

For optimizing the model in the case of obstacles, we need to perform this procedure in a 3D parameter space. We observed that the the parameter *ω* that models the response to the obstacles is only sensitive to changes on the order of 0.1. Since it is computationally intractable to sweep through the 3D parameter space in finer detail, we only look for values of *ω* ∈ [0, 1] in increments of 0.1, while keeping the same resolution for the other two parameters as done for the case of without obstacles. Furthermore, we found out that the two metrics *X* and *U* are not enough to obtain an optimal parameter set. However, this is expected as the increase in the degrees of freedom of the model may necessitate that an extra constraint must be satisfied. It is only natural to assume that the flow rate of the bats are likely the same with and without obstacles when they first exit the culvert, but when they reach the first line of obstacles, they are filtered such that not all of the bats will make it through. This is effect is evident when comparing Figures S1 C and D, where the lag at which the number of bats enter the control volume is slower in the case of obstacles. Therefore, another parameter that should account for the effect of bats’ response to the obstacles is the the distribution of entrance lag *L* = {*l*_1_, …, *l*_*N*−1_}, where *l*_*i*_ is number of timesteps of lag between the entrance of bats *i* and *i* − 1 into the control volume.

Figure 6 shows the heatmaps of energy distance for the metrics of *X* and *U* for the case without obstacles. The smallest threshold for which an intersection occurs is *δ* = 0.04, shown in the bottom plot. The centroid of this optimal intersection space is found to be {*χ* = 0.035, *σ* = 0.04}. For the case with obstacles, in Figure 7, we look for the optimal location in the heatmaps of energy distance for the metrics *X, U*, and *L*. The threshold for the case of obstacles is *δ* = 0.07 (please see Figure S2 for tuning over various *ω* values), and the centroid of the intersection space is found to be {*χ* = 0.009, *σ* = 0.015, *ω* = 0.2}. Table 1 shows a summary of these results.

**Fig. 7.**
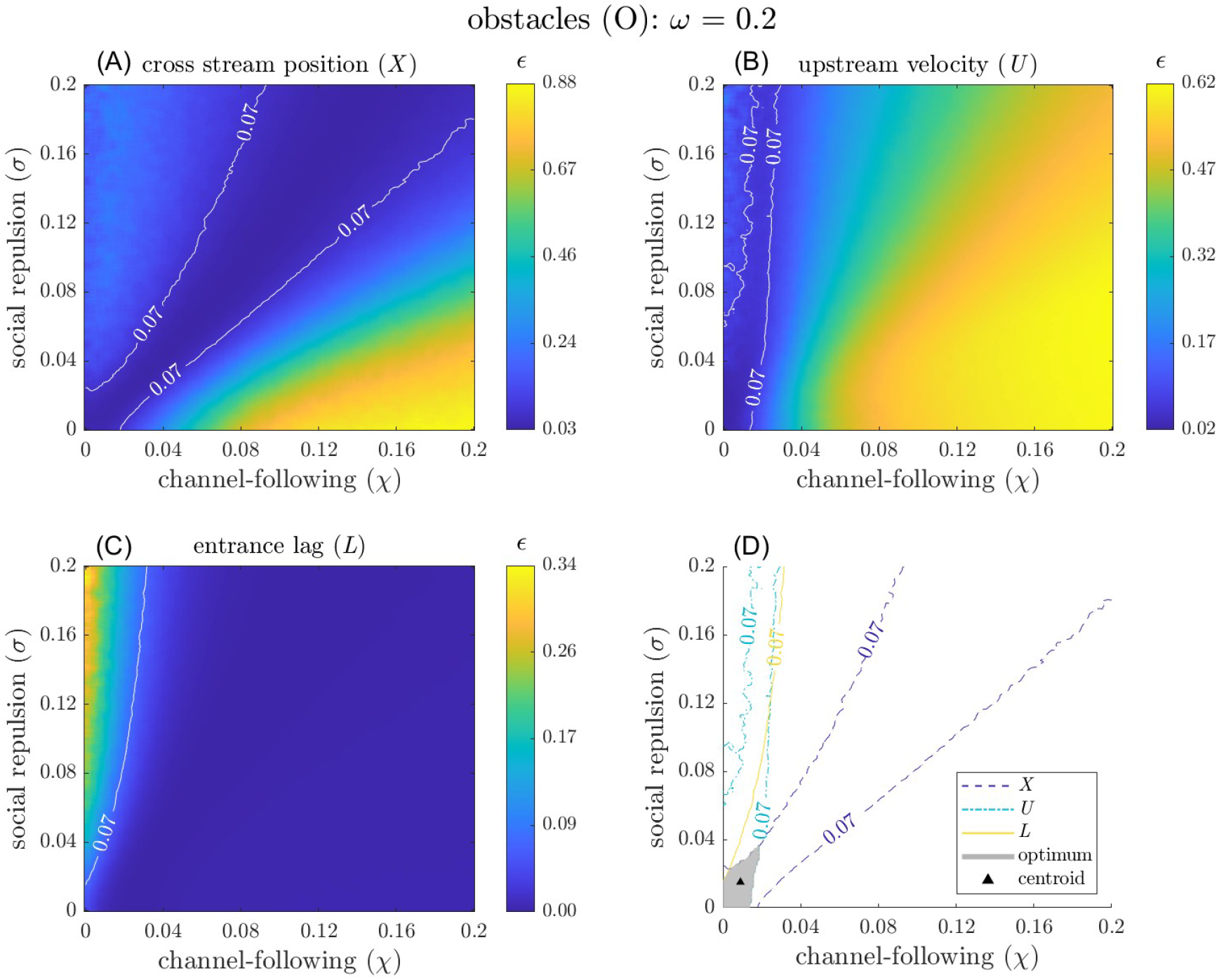
Heatmaps display values of energy distance, *ϵ*, computed using distributions of (A) cross stream positions, (B) upstream velocity, and (C) entrance lag. The color bars for each of the heatmaps represent the energy distances that are min-max normalized over the entire range of possible parameter values, i.e., *σ, χ, ω* ∈ [0, 1]. The heatmaps shown only consider the extent at which the optimal value is found, i.e., *ω* = 0.2 for obstacle avoidance, *χ* ∈ [0, 0.2] for channel-following, and *σ* ∈ [0, 0.2] for social repulsion. White lines overlaid on top of each heatmap represent the error threshold, *λ* = 0.07. (D) Energy distance contours with intersection in the parameter space depicted by the shaded gray region. The centroid of the optimum parameter set is shown by a black triangle at *χ* = 0.009 and *σ* = 0.015.

**Fig. 8.**
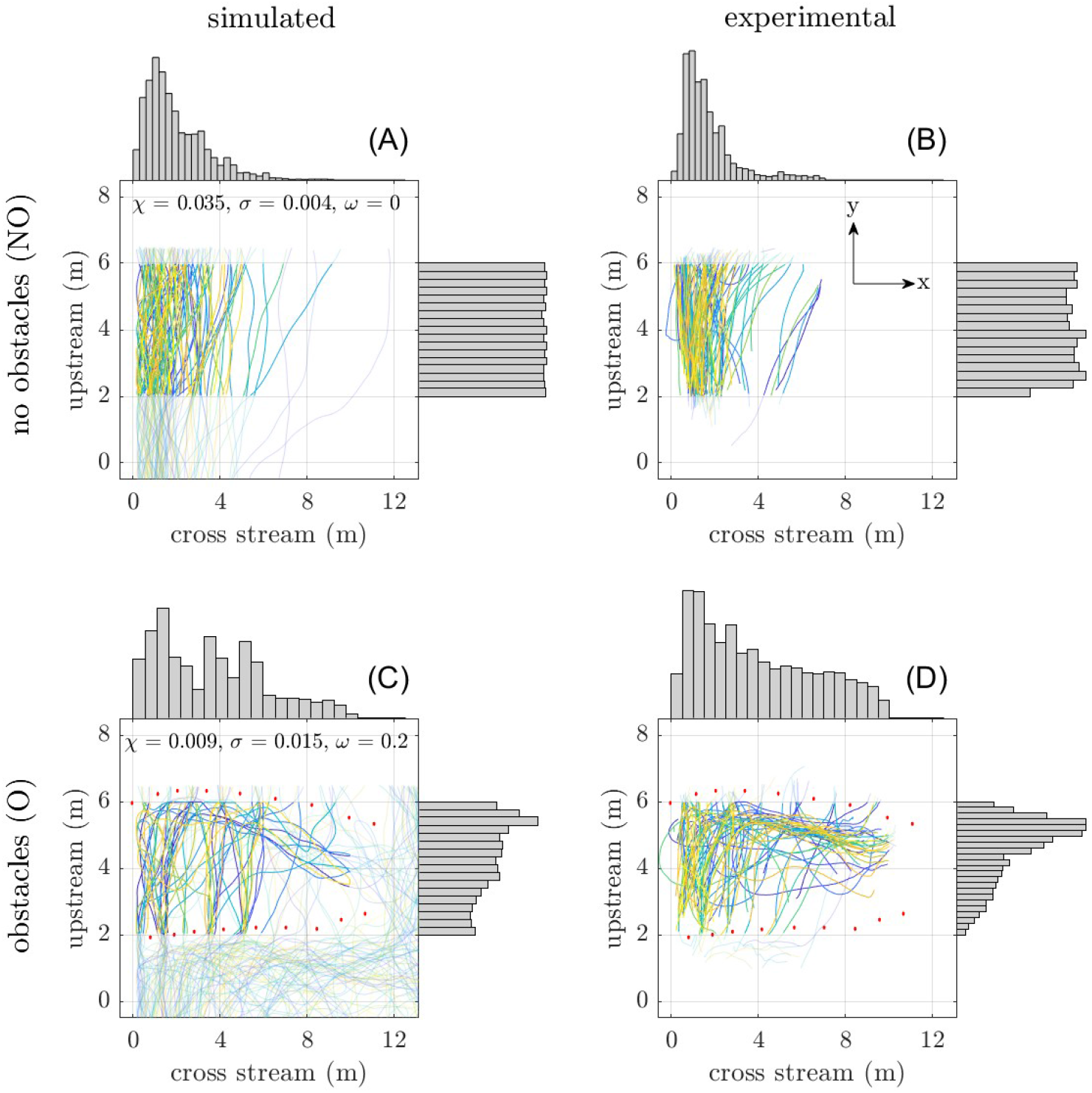
Comparison of simulated (left column) and experimental (right column) trajectories: The first row corresponds to the case of no obstacles (NO), and the second row corresponds to the case of obstacles (O). The values of *χ, σ*, and *ω* for the simulated trajectories are shown as insets, taken from the optimal parameter set in Table 1. Each of the colored lines shows the trajectory of a unique bat. The histograms on the top and right of each plot corresponds to the distribution of all the spatial points in their respective projected coordinate. The red circles on the bottom figures show the obstacle center locations.

## Discussion

In this study, we investigate how bats behave differently when they are challenged to navigate novel obstacles in their familiar flight paths. We learn that bats may have to renegotiate the balance of their social responses with interactions with the physical world. In our study, when faced with novel obstacles, bats shift from relying on wall-following to socially responsive and avoidance behaviors.

When navigating familiar flight paths, bats exhibit wall-following behavior but may spread out as a group that is likely modulated by social interactions. The spatial distribution of bats’ flight across the stream in Figure 4 (A) shows that, when flying alone, bats tend to stay near the left side of the channel and fly parallel to the left wall. The tendency to follow a wall or parallel structure has been observed in various animal species. This behavior may be attributed to the reliance on the parallel structure for navigational sensing, as observed in mice (35) and bees (36). Alternatively, it can reflect a stress response as seen in zebra fish (37). Both explanations could account for the behaviors we observe in singleton bats, since they could be relying on the walls as acoustic landmarks for navigation or they may stressed as social animals flying alone. Additionally, their choice to fly on the left side of the channel could be due to the presence of illumination from street lights, where bats are known to also rely on visual cues (38, 39). On the other hand, we observe an adjustment to this behavior when we observe their flights with contemporaneous neighbors. Our statistical analysis reveals that bats flying in groups tend to spread out across the stream, reflecting possible social interactions or strategies to reduce acoustic clutter which take priority over preferences for individual flight (24).

In the presence of obstacles, bats exhibit planar avoidance (or exploratory) turning flight patterns that we don’t see in their absence. We observe turning patterns even in the absence of sociality in Figure 4 (B). This suggests that bats may change their stereotypic flight paths when navigating a new (cluttered) environment, a behavior studied in big brown bats (*Eptesicus fuscus*) (16). Our statistical analysis further reveals that bats’ response to obstacles is planar as indicated by the significant change in velocity distributions up and across the stream but not in the direction of gravity. Interestingly, the planar response to vertical obstacles of the gray bats in our experiment is different to what is observed for big brown bats by Sandig *et al*. (12), which show that bat response is predominantly in the vertical direction. However, the vertical flight response to vertical obstacles in the case of the Sandig *et al*. also could be attributed to the presence of a landing platform along with prior training. Comparing Figures S1 and (B), we observe that the median speed is reduced when obstacles are present. This is consistent with previous findings in big brown bats (12), where slowing down allows more time for echolocation. When comparing singletons to groups, we find a significant difference in their upstream velocities only when obstacles are present. This suggests that turning behaviors that are socially driven may be more pronounced when bats are challenged by environmental perturbations.

Our agent-based model reveals that bats exhibit more stereotypy in familiar environments and shift to socially responsive and avoidance strategies when the environment has novel features that must be navigated. The model simulates channel-following as a proxy for their stereotypic behavior, and avoidance interactions to neighbors and obstacles as adaptations to the dynamic environment. By fitting the simulated trajectories to experimental data, we identify weights for each of the flight behaviors. In Figure 8 (A) and (B), we see that the optimized model captures both the normally distributed cross-stream and uniformly distributed upstream trajectories in the absence of obstacles. Additionally, when obstacles are introduced, Figure 8 (C) and (D), the shift toward normally distributed upstream trajectories is also observed in the optimized model. From comparing the optimized behavior weights, we find that channel-following behavior dominates social repulsion in the absence of obstacles by an order of magnitude difference. This behavioral state is flipped in the presence of obstacles, where the model shows that the weight of the social repulsion rule is an order of magnitude higher. This interesting finding indicates that social interactions become more pronounced when the environment becomes more challenging, where bats may spread out to avoid acoustic clutter.

The modeling results also highlight the interaction ruling that bats exhibit in the presence of environmental obstacles. Our statistical analysis only shows significance in a distance-based interaction, like attraction/repulsion, instead of velocity-based interaction, like alignment (please see Figure 5). The agreement between the model and the experimental data further supports the idea that, in the presence of environmental obstacles, bats’ primary objective is to mitigate acoustic clutter or simply avoid nearby conspecifics, in contrast to potentially more complex behaviors they may execute in less complicated environments. Our finding is in contrast to flocking behaviors, such as chasing interactions studied by Giuggioli *et al*. (26), which are more prominent in foraging or open-space scenarios.

In conclusion, our agent-based model informed by experimental data reveals that bats renegotiate their navigational behaviors depending on qualities of their environment. They tend to stay close to walls and follow the channel direction, but when introduced to novel obstacles, they show social behaviors in the form of spatial segregation. This indicates that bats can socially adapt their flight patterns in environmental clutter.

## ACKNOWLEDGEMENTS

We want to thank Megan Grey for her help assisting with data acquisition, and Dr. W. Mark Ford for helping with identifying a field site and obtaining permission to work there. We also want to thank Mihir Patel, Mariah Bolden, and Carolyne Bryant for their help in tracking bat trajectories. We want to acknowledge Virginia Tech’s Advanced Research Computing (ARC) for computing resources, and especially Dr. Matthew Brown and Dr. Chris Kuhlman for the computational advice.

## ETHICS

This study followed ethical standards set by the Virginia Tech Institutional Animal Care and Use Committee (IACUC 21-087) and was approved by the Virginia Department of Game and Inland Fisheries.

## DATA ACCESSIBILITY

The processed bat trajectories and the MATLAB scripts for the model are available at https://github.com/eighdiaung/ Agent-based-modeling-of-gray-bat-flight.git.

## AUTHORS’ CONTRIBUTIONS

Conceptualization: E.A. and N.A., Data curation: E.A., Formal analysis: E.A. and N.A, Funding acquisition: N.A., Methodology: E.A. and N.A., Software: E.A., Visualization: E.A., Writing - original draft: E.A., Writing - review and editing: E.A. and N.A. All authors reviewed the manuscript.

## Supplementary Section 1: Details on social interaction rules

To infer the presence of attraction or repulsion interactions, we fit a linear mixed effect model of the form:

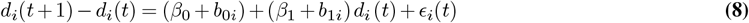

where *d*_*i*_(*t*) is the distance between bat *i* and its nearest neighbor at time *t*. For the linear model, *β*_0_ is the coefficient for the intercept, *b*_0*i*_ is the random effect for the intercept, *β*_1_ is the coefficient for the slope, *b*_1*i*_ is the random effect for the slope, and *E*_*i*_(*t*) is stochastic noise which is IID for all *i* and *t*. To infer the presence of flocking interactions, we fit the linear mixed effect model of the form:

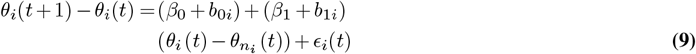

where *θ*_*i*_(*t*) is the heading of a bat *i* at time *t*, 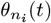 is the heading of the nearest neighbor bat, *β*_0_ is the coefficient for the intercept, *b*_0*i*_ is the random effect for the intercept, *β*_1_ is the coefficient for the slope, *b*_1*i*_ is the random effect for the slope, and *ϵ*_*i*_(*t*) is stochastic noise which is IID for all *i* and *t*. We note that, the independent and dependent variables are both normalized using the min-max norm for fair comparison across interaction types and experimental conditions.

## Supplementary Section 2: Details on model parameter identification

Some parameters can be directly estimated by computing the median values of their respective distributions we obtain from the data. The speed, *s*, at which bats fly is found to be 7 m/s for NO (Figure S1 A) and 5 m/s for O (Figure S1 B). We fix the time step, *τ* to be 1/60 s, which is equivalent to the sample rate of the video data. The time lag between the spawning of new agents into the simulation is found by fitting an exponential distribution to the entrance lags between any consecutive bats that enter the image area. We estimated *λ* = 0.24 s without obstacles (Figure S1 C) and *λ* = 0.35 s with obstacles (Figure S1 D). Although *λ* is indentified for the case with obstacles, we only use *λ* = 0.24 s when simulating the model since the flow rate of bats entering the experimental volume should be the same regardless. To characterize the behaviors for the bats’ interaction with geometry in the environment such as the wall and obstacles, we use the dataset from singleton bats to remove the possible effects of conspecifics. We find *R*_*w*_ = 0.8 m (Figure S1 E) and *R*_*o*_ = 0.5 m (Figure S1 F). The distance for social interactions is found from dataset when groups of bats are flying in the absence of obstacles, to remove the possible influence of obstacles on the social distancing. We find the minimum distance to the nearest neighbor bat is *R*_*s*_ = 1.1 m (Figure S1 G). The variance of the stochastic noise is fixed to be *η* = 0.05, from qualitative inspection of the simulations.

**Fig. S1.**
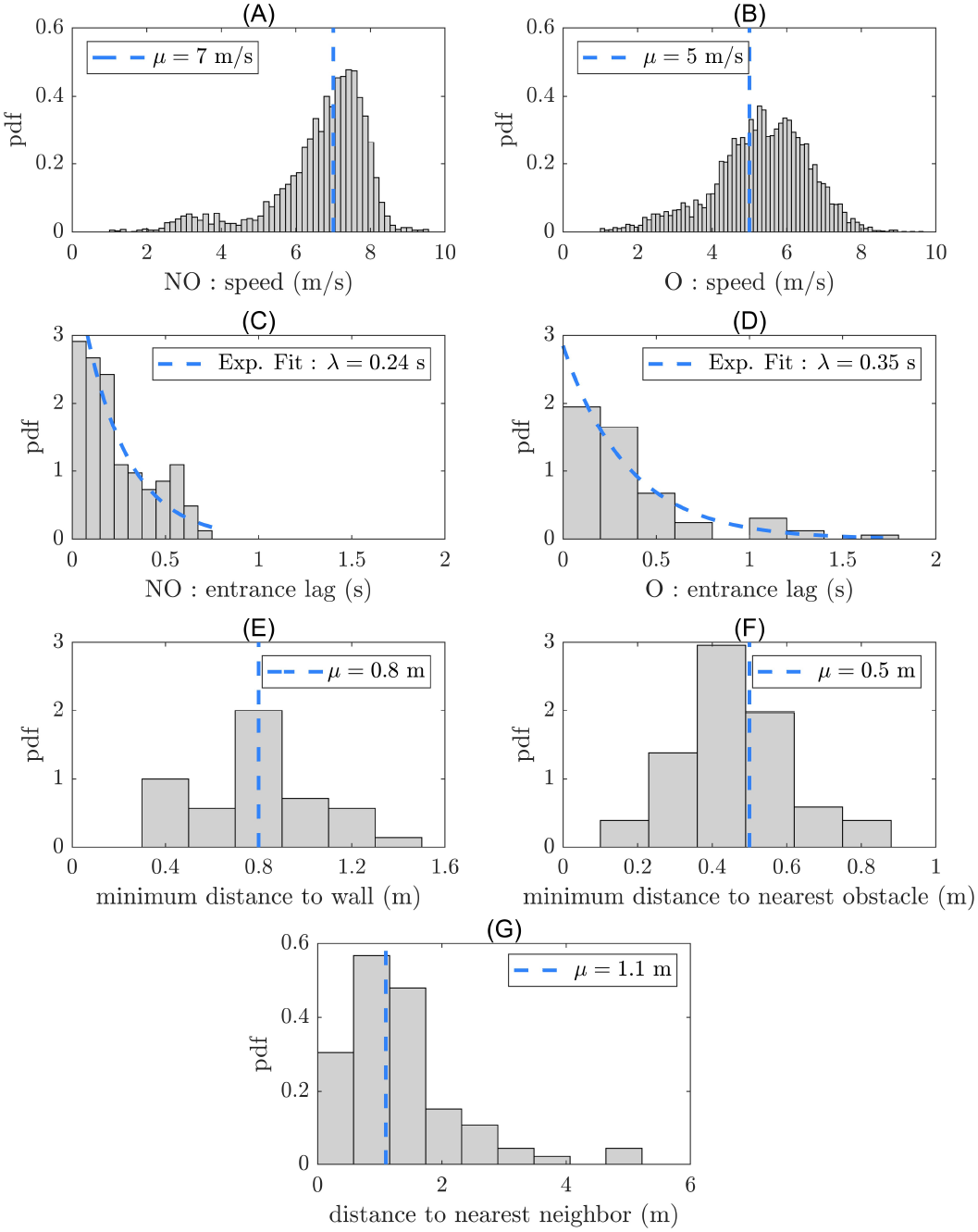
Model parameters obtained from experimental data: (A) Bats fly at median speed of 7 m/s when there are no obstacles (NO), and (B) at median speed of 5 m/s when obstacles are present (O), showing a slight decrease in median speed due to turning behavior in the presence of obstacles. The median time lag between the entrance of bats into the image area is 14 frames (~ 0.25 s) (C) when there are no obstacles, and (D) every 19 frames (~ 0.3 s) when there are obstacles. (E) Singleton bats fly at a median minimum distance of 0.8 m from the wall when there are no obstacles, and fly at a median minimum distance of 0.4 m from the obstacles. (G) When bats are flying in groups, the median distance to their nearest neighbor is 1.3 m (~ 5 wingspans).

**Fig. S2.**
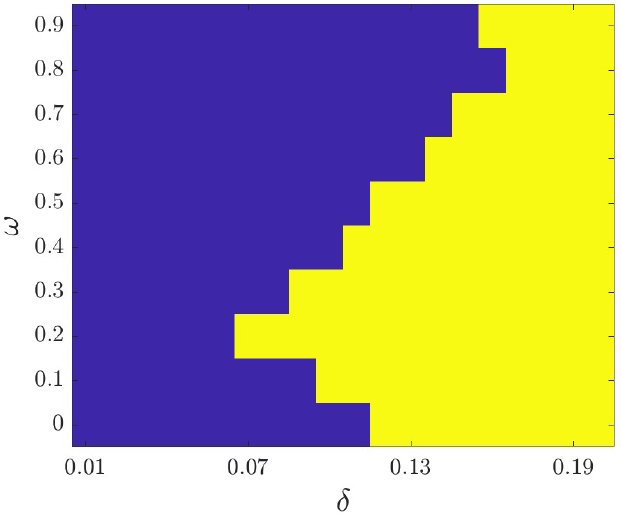
Parameter sweep for model with obstacles to find the minimum threshold, *δ*, where an intersection between metrics *X, U*, and *L* can be found for *ω* = [0, 9]. *δ* is tuned in increments of 0.01, and yellow/blue represents the existence/nonexistence of an intersection, respectively. The minimum *δ* is observed to be 0.07.

